# Along-Tract Microstructural Alterations Associated with Stimulant Misuse Localized using Diffusion MRI Tractometry

**DOI:** 10.64898/2026.03.28.714553

**Authors:** Leila Nabulsi, Yixue Feng, Bramsh Q. Chandio, Julio E. Villalon-Reina, Iyad Ba Gari, Jonathan Davis Alibrando, Talia M. Nir, Anthony C. Juliano, Devarshi Pancholi, Gwenyth S. Roundy, Nicola Canessa, Eduardo A. Garza-Villarreal, Hugh Garavan, Neda Jahanshad, Paul M. Thompson

## Abstract

Diffusion brain MRI (dMRI) studies of substance use disorders have reported widespread but modest white matter (WM) microstructural alterations with limited anatomical specificity. Here, we applied segment-wise along-tract 3D tractometry to brain dMRI scans to localize fine-scale WM alterations associated with stimulant misuse using two complementary analytical frameworks: Bundle Analytics (BUAN) and Medial Tractography Analysis (MeTA). We analyzed 3D profiles of widely-used diffusion metrics across 33 major WM bundles in independent cohorts of cocaine (74 cases;58 controls) and amphetamine (22 cases;18 controls) users, testing the statistical associations with brain microstructure of pooled stimulant effects, substance-specific effects, and direct comparisons between stimulant classes. Segment-wise analyses revealed focal differences localized to specific tract segments rather than uniform differences along entire bundles. In pooled stimulant misuse, convergent findings across analysis pipelines were localized to hippocampal pathways and were consistent with altered microstructural organization. Amphetamines misuse showed a broader pattern of segment-wise differences across commissural, projection, and association pathways, involving altered axonal organization. No robust segment-wise differences were detected for cocaine misuse or between stimulant classes. These results show that WM alterations are spatially localized and reproducible across tractometry frameworks, highlighting the value of along-tract 3D mapping for improving anatomical specificity in addiction neuroimaging.

## I. Introduction

Substance misuse disorders are associated with persistent neurobiological alterations affecting large-scale brain circuits that support reward learning, executive control, and affect regulation [1– 3]. In neuroimaging studies of addiction, a key question is whether brain abnormalities differ depending on the substance used (e.g., methamphetamine versus alcohol versus cannabis), and if there are regionally specific or circuit specific abnormalities that either result from, or predispose people to, substance misuse. Among several neuroimaging techniques used to study addiction, and the effect of substance misuse in the brain, diffusion-weighted magnetic resonance imaging (dMRI) has been widely used to investigate white matter (WM) microstructure, providing indirect markers of axonal organization and myelination through metrics such as fractional anisotropy (FA) and mean diffusivity (MD). Diffusion MRI is sensitive to tissue abnormalities not detectable with standard anatomical MRI, and can also be used for fiber tracking, where macroscopic neural pathways can be reconstructed that represent the major fiber bundles of the brain. Across stimulant use disorders, prior dMRI studies report consistent abnormalities, at the group level, in fronto-striatal, limbic, and commissural pathways, implicating white matter circuitry as a key substrate of addiction-related pathology [1–3]. Large-scale, harmonized analyses, that integrate data across multiple sites and cohorts, have improved the reproducibility and interpretability of dMRI findings in substance misuse. In particular, the ENIGMA-Addiction consortium has applied tract-based spatial statistics (TBSS) to evaluate WM microstructural alterations across multiple substance use disorders in ∼800 participants worldwide [4]. These studies show that stimulant misuse is associated with widespread but non-uniform WM alterations, with prominent effects in limbic and interhemispheric pathways and more variable involvement of association and projection systems. Such findings provide compelling evidence that addiction-related WM alterations are distributed at the network level rather than confined to isolated regions. However, TBSS and related region-of-interest or skeleton-based approaches summarize diffusion properties over relatively large regions of the WM. This coarse spatial scale may obscure anatomically localized or more concentrated effects along individual fiber bundles. Averaging diffusion metrics across large regions or projecting them onto a common skeleton may also reduce sensitivity to focal abnormalities, particularly if effects vary along the length of a tract or are confined to specific subsegments. This limitation is arguably relevant in substance misuse, as neurotoxic exposure, compensatory plasticity, and circuit-specific vulnerability may manifest heterogeneously across WM pathways [5,6]. Along-tract tractometry has emerged as a complementary framework to region-of-interest analyses, that spatially resolves diffusion properties in 3D, along anatomically defined fiber bundles. By modeling diffusion metrics as functions of position along a tract, tractometry allows for detection of focal microstructural alterations that may not be apparent in region-of-interest or skeleton-based analyses [7–9]. Within this framework, multiple implementations have been proposed, including BUndle ANalytics (BUAN), which characterizes segment-wise diffusion profiles along population-derived fiber bundles [8], and medial-curve–based parameterization approaches such as Medial Tractography Analysis (MeTA) [9]. Both methods sample diffusion properties along a tract’s medial trajectory to estimate a computed anatomical correspondence across subjects, and enhance statistical sensitivity. A range of tractometry and bundle-based mapping methods have been developed to support anatomically informed mapping of WM microstructure, including Automated Fiber Quantification (AFQ) [7], TRActs Constrained by UnderLying Anatomy (TRACULA) [10], TractSeg [11], and more recently, related approaches detailed in [12]. In the present work, we focus on BUAN and MeTA as complementary tractometry methods, although a more exhaustive analysis may be worthwhile in future. Despite these advances, along-tract methods have not yet been applied in large-scale addiction studies, and the robustness of segment-wise findings across independent tractometry frameworks has not been examined. In the present study, we create 3D maps of WM alterations associated with stimulant misuse using two independent cohorts of cocaine and amphetamine users; this pilot study is intended to motivate a larger scale, multicohort study. We apply two complementary tractometry frameworks, BUAN and MeTA, to analyze diffusion profiles along 33 major WM bundles in the brain. Rather than treat these approaches as competing methods, we use them as largely independent analytic approaches to assess the consistency and robustness of observed effects. We test three contrasts of interest: (i) all stimulant misuse (cocaine and amphetamine combined) versus non-users, (ii) substance-specific effects comparing cocaine users and amphetamine users to their respective control groups, and (iii) direct comparisons between cocaine and amphetamine users. Our primary objective is to determine whether focal, segment-wise WM alterations associated with stimulant misuse are reproducibly identified using independent tractometry frameworks, and to identify the functional systems involved. We hypothesize that stimulant misuse is associated with focal microstructural alterations preferentially affecting limbic and fronto–striatal/thalamocortical pathways, consistent with established models of addiction circuitry [1-3, 13], and that amphetamine misuse may exhibit a more spatially distributed pattern of effects relative to cocaine misuse. By extending prior ENIGMA-Addiction TBSS findings toward anatomically precise, spatiallyt-resolved analyses, this work aims to refine the characterization of WM alterations in stimulant misuse and to understand the potential utility of tractometry for circuit-level investigations for large, multi-site neuroimaging studies.

## II. Methods

### A. Participants & Study Sample

We analyzed data from two independent cohorts of stimulant users recruited at separate sites, comprising individuals with amphetamine misuse and individuals with cocaine misuse, along with demographically matched non-user control groups at each site. All participants provided informed consent in accordance with local institutional review board approvals. The cocaine misuse cohort consisted of 74 cases (mean age ± SD: 31.0±7.6 years; 87.8% males) and 58 control participants (30.8±7.5 years; 81.0% males). The amphetamine misuse cohort comprised 22 cases (45.6 ± 8.0 years; 59.1% males) and 18 control participants (44.8 ± 8.9 years; 61.1% males). Control participants were recruited at each site and matched for age and sex. In addition to substance-specific analyses, a combined ‘all stimulant misuse’ sample was constructed by pooling amphetamine and cocaine misuse cases and their respective controls. This combined sample was used to examine shared WM alterations associated with stimulant misuse independent of substance type, while substance-specific analyses were performed within each cohort to preserve site-specific and demographic structure.

### B. Diffusion MRI Processing & Statistical Analyses

All diffusion MRI data were preprocessed using PreQual [14], an automated pipeline for integrated preprocessing and quality assurance of diffusion-weighted imaging data. PreQual performs denoising, Gibbs ringing correction, correction for susceptibility-induced distortions, eddy-current–induced distortions, and subject motion, with appropriate rotation of diffusion gradient directions. Brain extraction was applied to generate binary diffusion masks used in downstream analyses. PreQual also generates quantitative and visual quality control (QC) outputs to help in excluding scans affected by excessive motion or artifacts. In the resulting preprocessed diffusion images, cerebellar regions frequently exhibited visible artifacts and insufficient data quality, precluding reliable tract reconstruction in that region. As a result, cerebellar bundles were not included in the final analyses. Whole-brain diffusion information was used to perform bundle-specific probabilistic tractography guided by a modified version of the population-averaged HCP1065 WM atlas [15]. Fiber orientation distributions were calculated using constrained spherical deconvolution [16] and tractography was performed separately for each major WM bundle. Subject DWI data were nonlinearly registered to the MNI space using multichannel ANTs Syn [17], where FA, MD and the mean b0 image were jointly used to estimate the deformation field. Atlas bundles transformed to the subject space were used to derive seeding and stopping regions. Probabilistic tracking was performed with a fixed step size of 0.2 mm, and a bundle-specific maximum turning angle ranging from 20 to 30°. Streamline lengths were constrained using a relative length tolerance of ± 20% with respect to the corresponding atlas bundle. Fast Streamline Search was used to further remove implausible streamlines [18]. Additional constraints included an endpoint filtering tolerance of 3 mm, dilation of seeding and stopping masks to improve spatial coverage, and enforcement of a minimum of 20 streamlines per bundle. Bundles that failed to meet streamline count or quality criteria were excluded from downstream analyses. Reconstructed bundles were next spatially normalized to standard MNI space using the precomputed ANTs transform. Registration quality was assessed by visually inspecting the alignment between diffusion images and the template. QC of bundle reconstructions was performed at both the subject and bundle levels. Visual inspection of reconstructed bundles was conducted to identify gross anatomical inconsistencies or tracking failures. In addition, automated QC screening was applied to identify outlier subjects or bundles based on predefined thresholds. Bundles failing QC were excluded from tractometry analyses. Segment-wise WM microstructural profiles were extracted using two independent along-tract tractometry frameworks: streamline-based Bundle Analytics (BUAN) [8] and voxel-based Medial Tractography Analysis (MeTA) [9], both depicted in **Fig. 1**. Both methods were applied to the same set of reconstructed WM bundles and diffusion maps, enabling complementary characterization of focal microstructural variation along tract trajectories.

**Figure 1.**
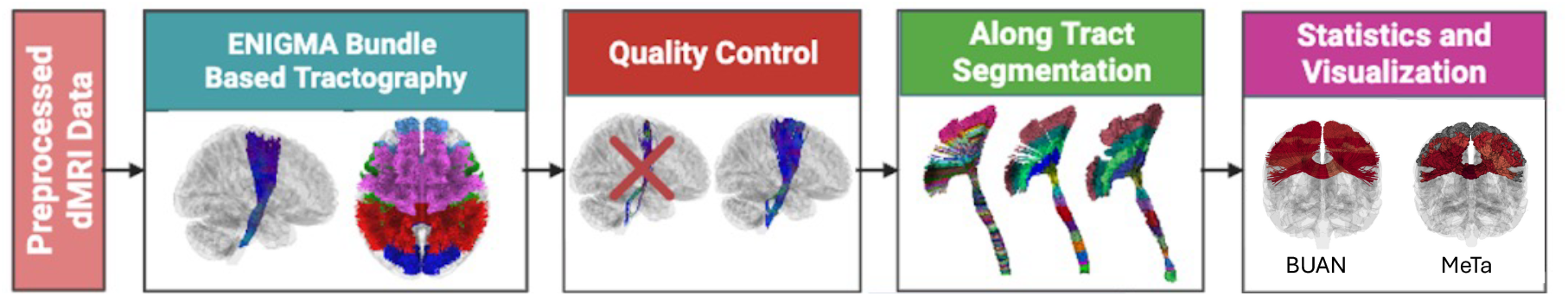
Overview of the Tractometry processing pipeline for BUAN and MeTa. Preprocessed diffusion MRI data undergo bundle-based tractography, quality control, along-tract segmentation, and segment-wise statistical analysis tractometry frameworks.

### C. Bundle ANalytics (BUAN)

BUAN tractometry was used to extract segment-wise along-tract diffusion profiles along each reconstructed WM bundle. For each subject and bundle, streamlines were parameterized relative to the atlas bundle to ensure anatomical correspondence across individuals. Diffusion metrics from diffusion tensor imaging (DTI) including FA, MD, radial diffusivity (RD), and axial diffusivity (AxD), were sampled along the tract trajectory and summarized at fixed-length segments. Sampled diffusion metrics were averaged within each segment weighted by distance to the bundle centroid to generate a one-dimensional tract profile per subject [19]. This procedure yielded anatomically aligned, segment-wise diffusion profiles suitable for statistical modeling across subjects.

### D. Medial Tractography Analysis (MeTA)

MeTA tractometry was used to derive complementary segment-wise profiles based on medial-curve parameterization of WM bundles. For each subject and bundle, streamlines were first converted into a volumetric representation from which a medial surface was estimated. The medial trajectory representing the core 25% of the bundle was then computed, providing a compact and anatomically stable representation of the tract. Bundles were subdivided into a predefined number of segments along the medial trajectory [20], with segment definitions consistent across subjects. Diffusion metrics (FA, MD, RD, and AD) were sampled within each medial segment from subject-specific diffusion maps, producing segment-wise volumetric profiles. This approach emphasizes anatomical correspondence by anchoring measurements to the bundle’s medial core rather than to individual streamline trajectories. Segment-level diffusion profiles were extracted independently for each bundle and subject.

### E. Statistical analysis

Segment-wise statistical analyses were conducted on along-tract diffusion profiles derived from both tractometry frameworks. Segment-wise linear mixed-effects models (LMMs) were used to assess group differences in diffusion metrics, with age at MRI scan and sex included as covariates. Site was modeled as a random intercept when analyses involved data from more than one acquisition site. Three contrasts were examined: (i) All stimulant misuse versus non-users, assessed by pooling amphetamine and cocaine cohorts and modeling group status as the effect of interest; (ii) Substance-specific effects, assessed separately within amphetamine and cocaine cohorts by comparing misuse cases to matched controls, without modeling site as a random effect due to single-site analyses; and (iii) Between-substance differences, assessed by directly comparing cocaine and amphetamine misuse cases, with site included as a random effect when applicable. Several WM bundles were excluded following bundle-level QC due to poor reconstruction reliability, primarily involving cerebellar and inferior brainstem pathways. To ensure stable model estimation, analyses were restricted to bundles meeting minimum sample size requirements. Bundles were excluded if fewer than 30 subjects were available overall. Bundles or segments for which model fitting failed or returned empty results were excluded from further analysis. Multiple-comparison correction was applied using the Benjamini– Hochberg false discovery rate (FDR) [21] across all bundles within each diffusion metric (α<0.05). Summary statistics were generated at both the segment and bundle levels to help in interpreting and visualizing focal WM effects [22].

## III. Results

Across contrasts of interest, reported findings were localized to discrete segments within specific WM bundles and were consistent across tractometry frameworks for the tracts and metrics. Results are reported only for tract–metric combinations showing FDR-corrected significance and representing the strongest and most spatially extensive effects, as defined by the highest number of significant segments across both tractometry frameworks, and are depicted in **Fig. 2**. For each depicted tract and diffusion metric, summary statistics, including the minimum FDR-corrected p-value, the number and proportion of significant segments, and the mean segment-wise effect size expressed as partial Cohen’s *d*, are provided in **Tab 1**. In the pooled stimulant misuse contrast, significant segment-wise effects were identified within hippocampal pathways using both tractometry frameworks. BUAN analyses identified lower FA in localized regions of the left hippocampal alveus. MeTA analyses identified lower FA, and higher MD and RD, in localized segments of the right hippocampal alveus. In the amphetamine-specific contrast, segment-wise effects were identified across multiple WM bundles for BUAN and MeTA, with lower FA seen in localized segments of the left anterior corticospinal pathway, higher MD in posterior callosal fibers (tapetum), and both higher MD and RD in the anterior body of the corpus callosum. Higher AxD was seen in localized regions of the right anterior radiation. No segment-wise effects meeting robustness criteria were identified in the cocaine misuse versus control contrast or in the direct comparison between amphetamine and cocaine users.

**Figure 2.**
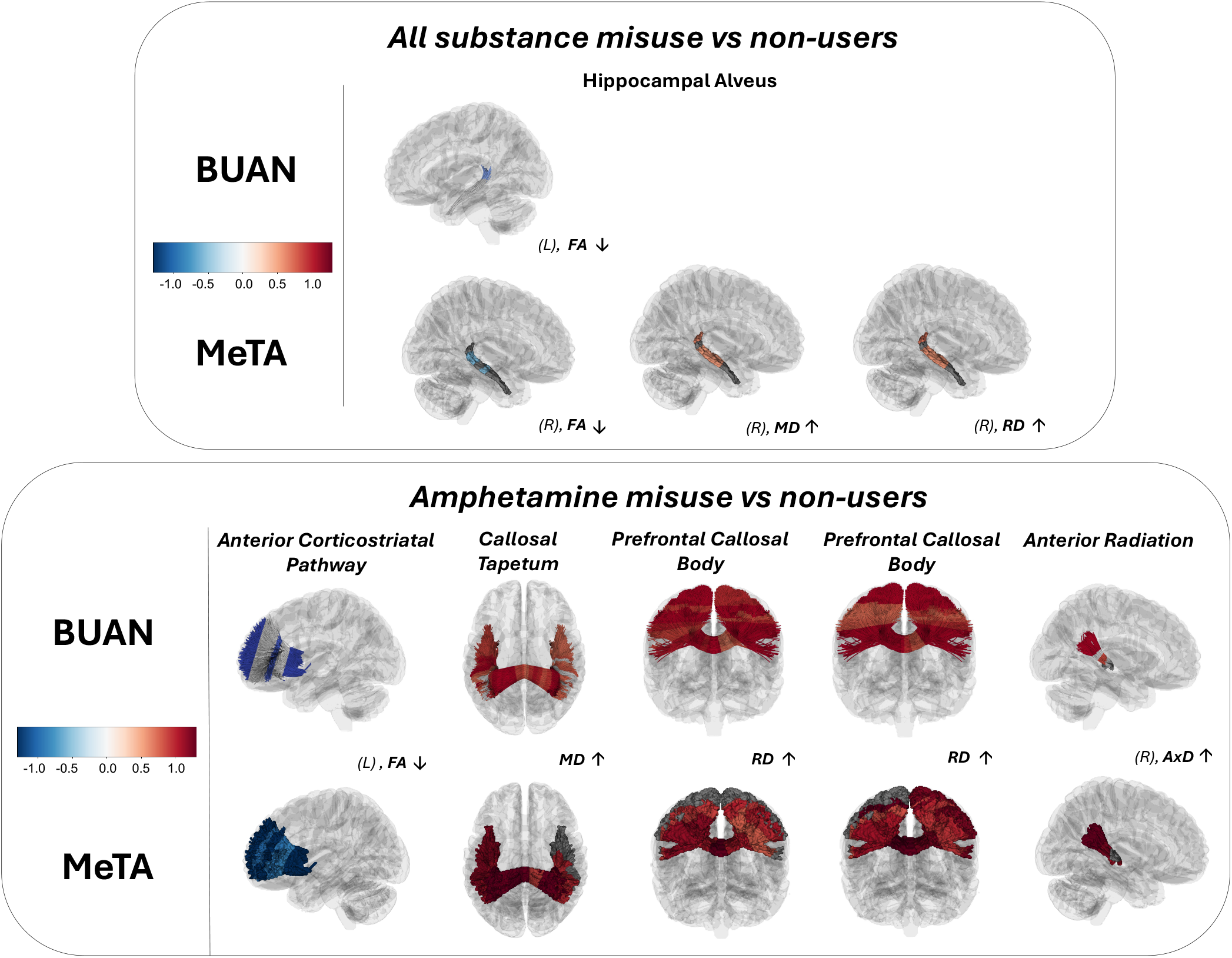
Segment-wise group differences in white matter microstructure. Results are shown for representative tracts across two contrasts: all substance misuse (cocaine + amphetamine) vs non-users (top panel) and amphetamine misuse vs non-users (bottom panel). Results are displayed for multiple white matter microstructural metrics, with color values representing partial Cohen’s *d* at each tract segment. Highlighted tract segments indicate locations exhibiting statistically significant group differences after false discovery rate correction (*p*FDR < 0.05). Across both analysis frameworks (BUAN and MeTA), effects in the all substance misuse contrast are confined to limbic pathways, whereas the amphetamine-specific contrast shows more distributed involvement of limbic, commissural, and fronto–striatal/thalamocortical projection systems. Effects are localized to specific tract segments rather than uniformly distributed along entire tracts. Tracts shown were selected based on consistency of effect directionality, functional relevance to addiction-related circuitry, and convergence methods. L = left-hemisphere; R = right-hemisphere; FA = fractional anisotropy; MD = mean diffusivity; RD = radial diffusivity; AxD = axial diffusivity.

**Table 1.** Convergent segment-wise white matter findings across tractometry frameworks. The table summarizes focal white matter alterations identified using along-tract tractometry and retained based on convergence across analytical frameworks, consistent anatomical localization, and consistent directionality. Results are reported for each contrast, tractometry framework (BUAN or MeTA), white matter bundle, and diffusion metric. *Min FDR-p* denotes the minimum false discovery rate–corrected *p*-value across significant segments within a bundle. *Sig segments* indicate the number of segments surviving FDR correction relative to the total number of segments tested, with the segment index range shown in parentheses. *FDR proportion* represents the fraction of segments for each bundle that survived FDR correction. *Mean partial d* corresponds to the mean segment-wise partial Cohen’s *d* across significant segments. Direction arrows indicate whether diffusion metric values were higher (↑) or lower (↓) in cases relative to controls.

| Contrast | Method | Direction vs controls | DWI Metric | Anatomical Region | Functional system | Min FDR-$p$ | Mean Partial $d$ | Sig segments n/N (range) | FDRproportion |
| --- | --- | --- | --- | --- | --- | --- | --- | --- | --- |
| All stimulants use vs non-users | BUAN | ↓ | FA | Hippocampal Alveus (Left) | Limbic | 0.037 | -0.6 | 2/10 (8, 9) | 0.2 |
|  | MeTA | ↓ | FA | Hippocampal Alveus (Right) | Limbic | 0.045 | -0.6 | 3/9 (4, 6, 7) | 0.3 |
|  | MeTA | ↑ | MD | Hippocampal Alveus (Right) | Limbic | 0.036 | 0.6 | 5/9 (3:6, 8) | 0.6 |
|  | MeTA | ↑ | RD | Hippocampal Alveus (Right) | Limbic | 0.035 | 0.6 | 5/9 (3:6, 8) | 0.6 |
| Amphetamines use vs non-users | BUAN | ↑ | AxD | Anterior Radiation (Right) | Projection | < 0.0001 | 1.6 | 5/12 (6, 8:11) | 0.4 |
|  | MeTA | ↑ | AxD | Anterior Radiation (Right) | Projection | 0.0003 | 1.3 | 4/7 (1, 4:6) | 0.6 |
|  | BUAN | ↑ | MD | Corpus Callosum (Anterior body) | Commissural | < 0.0001 | 1.3 | 24/25 (1:24) | 1.0 |
|  | MeTA | ↑ | MD | Corpus Callosum (Anterior body) | Commissural | < 0.0001 | 1.1 | 11/14 (2:12) | 0.8 |
|  | BUAN | ↑ | RD | Corpus Callosum (Anterior body) | Commissural | < 0.0001 | 1.2 | 24/25 (1:24) | 1.0 |
|  | MeTA | ↑ | RD | Corpus Callosum (Anterior body) | Commissural | < 0.0001 | 1.2 | 12/14 (2:13) | 0.9 |
|  | BUAN | ↑ | MD | Corpus Callosum (Tapetum) | Commissural | < 0.0001 | 1.2 | 30/35 (1:2, 4:6, 9:25, 27:34) | 0.9 |
|  | MeTA | ↑ | MD | Corpus Callosum (Tapetum) | Commissural | < 0.0001 | 1.4 | 16/18 (2:17) | 0.9 |
|  | BUAN | ↓ | FA | Anterior Corticospinal Pathway (Left) | Projection | < 0.0001 | -1.6 | 4/11 (1, 6, 9:10) | 0.4 |
|  | MeTA | ↓ | FA | Anterior Corticospinal Pathway (Left) | Projection | < 0.0001 | -1.4 | 7/8 (1:7) | 0.9 |

## IV. Discussion

In this study, we applied segment-wise tractometry to characterize localized WM alterations associated with stimulant misuse across two independent cohorts, using BUAN and MeTA as complementary analytical frameworks. By emphasizing convergence across methods rather than method-specific effects, we identified reproducible and anatomically localized alterations confined to specific tract segments and functional systems. This approach extends prior large-scale dMRI findings in addiction by refining the spatial specificity of reported effects. The ENIGMA Addiction Working Group previously reported widespread but modest WM alterations across substance use disorders using tract-based spatial statistics (TBSS), with effects most consistently observed in *corpus callosum, cingulum*, and projection pathways [4]. These findings were interpreted as reflecting shared vulnerability or chronic neurobiological adaptations associated with substance misuse, rather than substance-specific signatures. ENIGMA’s TBSS analysis also emphasized that effects were spatially heterogeneous across the regions included in the WM skeleton, motivating the use of complementary approaches that may provide greater anatomical resolution along individual fiber pathways. The present results are broadly consistent with this interpretation, while providing additional anatomical resolution. Rather than identifying diffuse alterations along entire tracts or skeleton regions, 3D tractometry revealed focal clusters of altered diffusion metrics confined to discrete portions of specific bundles, suggesting that effects observed in TBSS analyses may arise from localized microstructural alterations that are spatially averaged or may even be diluted in voxel-wise approaches. In the pooled stimulant misuse contrast, convergent effects across tractometry frameworks were restricted to hippocampal pathways, specifically the alveus. This localization is consistent with ENIGMA TBSS-based reports implicating limbic and paralimbic WM regions across substance use disorders [4], while refining their anatomical specificity to specific regions within hippocampal efferent pathways. These effects were characterized by lower fractional anisotropy alongside higher mean and radial diffusivity, consistent with altered microstructural organization and less restricted diffusion within these limbic fibers. As in prior work, such interpretations remain probabilistic, as diffusion metrics are sensitive to multiple, potentially co-occurring microstructural processes [23–25]. In contrast to the pooled analysis, amphetamine-specific comparisons revealed segment-wise alterations spanning commissural, projection, and association systems, including callosal pathways, anterior radiations, and corticospinal projections. This broader spatial distribution parallels ENIGMA TBSS observations that stimulant-related effects may extend beyond limbic regions to involve fronto-striatal and thalamocortical circuitry [4]. Segment-wise tractometry refines this picture as it shows that such involvement is not uniform along entire bundles but instead concentrated inspecific tract segments and associated with metric-specific directionality. Specifically, increases in mean and radial diffusivity were observed in callosal subregions, while projection pathways showed higher axial diffusivity and lower fractional anisotropy, suggesting distinct patterns of altered tissue diffusion constraints and axonal organization across pathway classes. These associations should not be interpreted mechanistically in isolation but as complementary indicators of microstructural remodeling [23-25]. 3D profiles of abnormality in amphetamine misuse were associated with moderate-to-large partial effect sizes, with partial Cohen’s *d* values exceeding those typically reported in large-scale voxel-wise TBSS analyses. This finding likely reflects the increased anatomical specificity afforded by along-tract approaches, which reduce spatial averaging and enhance sensitivity to focal alterations, rather than indicating stronger or more widespread biological effects. No robust segment-wise effects were identified in cocaine-specific analyses or in direct comparisons between amphetamine and cocaine users. This absence is consistent with ENIGMA TBSS findings, which reported limited substance-specific differentiation and substantial overlap in WM alterations across substance use disorders [4]. The lack of detectable cocaine-specific effects in the present study may reflect low power at the current sample size, greater heterogeneity, or lower spatial consistency of effects at the segment level. A key contribution of this work is the demonstration that independent tractometry frameworks generally converge on similar anatomical patterns, despite differences in parameterization strategy. This convergence suggests that the dentified effects are robust and that regional WM alterations associated with stimulant misuse can be reliably detected when analyses emphasize anatomical correspondence along tracts. This study does not aim to establish superiority of one tractometry method over another, but rather to demonstrate that convergent biological signals emerge across complementary analytical frameworks. Although the present study does not permit mechanistic inference, the anatomical distribution of findings may have relevance for functional domains consistently implicated in addiction. Limbic pathway involvement in the pooled stimulant misuse contrast aligns with the role of hippocampal circuitry in contextual learning, memory, and emotional processing, which are known to contribute to cue reactivity and relapse vulnerability [6]. In amphetamine misuse, the broader involvement of commissural and projection pathways encompasses systems supporting interhemispheric integration and fronto–subcortical communication, which are central to executive control, action selection, and behavioral regulation [6]. From this perspective, the observed localized WM alterations may reflect circuit-level adaptations or vulnerabilities in networks repeatedly implicated in stimulant misuse, rather than isolated or substance-specific pathology. Importantly, these interpretations remain descriptive and should not be construed as evidence of causal or molecular mechanisms. Several limitations should be acknowledged. Statistical power may have been insufficient to detect subtle or heterogeneous effects, particularly in smaller cohorts. The cross-sectional design precludes inferences about causality, premorbid vulnerability, or recovery effects. Although the direct comparison of substances did not yield significant effects, we note that the design is confounded as people using different substances were assessed at different scanners and sites, so any detected differences would have to be inferred in the light of these potential confounds. Diffusion MRI metrics provide indirect indices of microstructure and cannot uniquely map onto specific cellular or molecular mechanisms. Longitudinal and multimodal studies integrating local tractometry with complementary imaging measures (such as resting state functional MRI) or clinical measures will be required to further clarify the biological substrates and temporal dynamics underlying these focal alterations. In summary, 3D tractometry reveals focal, anatomically specific WM alterations associated with stimulant misuse that are consistent across independent analytical frameworks. These findings extend prior ENIGMA TBSS results by showing that WM differences in addiction can be resolved at a finer spatial scale using along-tract approaches. Our results show that 3D tractometry can help us to localize spatially localized alterations on WM bundles, and reveal spatial heterogeneity that may be not possible to detect by skeleton-based analyses. Together, these results highlight the value of along-tract approaches for improving anatomical specificity (spatial resolution and signal-to-noise) in large-scale neuroimaging studies of addiction, which is critical for mechanistic interpretation and for identifying features most relevant to disease processes.

## Acknowledgment

LN was supported by Brain & Behavior (NARSAD) Young Investigator Grant; LN, PMT and NJ were supported by R01AG057892; R01MH134004, RF1NS136995 (to PMT and NJ); we also acknowledge support from the “Ricerca Corrente” funding scheme of the Italian Ministry of Health to Istituti Clinici Scientifici Maugeri IRCCS to NC. EAGV received funding from CONACYT-FOSISS project No. 0201493, CONACYT-Cátedras project No. 2358948, Instituto de Neurobiología of the Universidad Nacional Autónoma de México (UNAM) and Programa de Anovo a Provectos de Investigacion e Innovacion Tecnologica (PAPIIT) DGAPA Grants IA202120 and IA201622. EAGV also received support from M. Mallar Chakravarty and Gabriel A. Devenyi at the Computational Brain Anatomy Lab (CoBrA Lab) (http://cobralab.ca/), CIC, Douglas Research Center, Montreal and Digital Research Alliance Canada (www.computecanada.ca) and Mallar Chakravarty (Director of the Computational Neuroanatomy Laboratory, Douglas Research Centre, Montreal, Canada) who provided access to computational tools from him group on the Niagara Compute Cluster. HG, AJ, DP were supported by R01DA047119.

## Notes

### Competing Interest Statement

The authors have declared no competing interest.

## References

[1]. Goldstein, R. Z., & Volkow, N. D. (2011). Dysfunction of the prefrontal cortex in addiction: neuroimaging findings and clinical implications. Nature Reviews Neuroscience, 12(11), 652–669. 10.1038/nrn3119

[2]. Koob, G. F., & Volkow, N. D. (2016). Neurobiology of addiction: a neurocircuitry analysis. The Lancet Psychiatry, 3(8), 760–773. 10.1016/S2215-0366(16)00104-8

[3]. Moeller, F. G., Hasan, K. M., Steinberg, J. L., et al. (2005). Reduced anterior corpus callosum white matter integrity is related to increased impulsivity and reduced discriminability in cocaine-dependent subjects. Neuropsychopharmacology, 30(3), 610–617. 10.1038/sj.npp.1300617

[4]. Ottino-González, J., Uhlmann, A., Hahn, S., Cao, Z., Cupertino, R. B., Schwab, N., Allgaier, N., Alia-Klein, N., Ekhtiari, H., Fouche, J.-P., Goldstein, R. Z., Li, C.-R., Lochner, C., London, E. D., Luijten, M., Masjoodi, S., Momenan, R., Oghabian, M. A., Roos, A., Stein, D. J., Stein, E. A., Veltman, D. J., Verdejo-García, A., Zhang, S., Zhao, M., Zhong, N., Jahanshad, N., Thompson, P. M., Conrod, P., Mackey, S., & Garavan, H. (2022). White matter microstructure differences in individuals with dependence on cocaine, methamphetamine, and nicotine: Findings from the ENIGMA-Addiction Working Group. Drug and Alcohol Dependence, 230, 109185. 10.1016/j.drugalcdep.2021.109185

[5]. Zalesky, A., Fornito, A., & Bullmore, E. T. (2010). Network-based statistic: identifying differences in brain networks. NeuroImage, 53(4), 1197–1207. 10.1016/j.neuroimage.2010.06.041

[6]. Fields, R. D. (2008). White matter in learning, cognition and psychiatric disorders. Trends in Neurosciences, 31(7), 361–370. 10.1016/j.tins.2008.04.001

[7]. Yeatman, J. D., Dougherty, R. F., Myall, N. J., Wandell, B. A., & Feldman, H. M. (2012). Tract profiles of white matter properties: automating fiber-tract quantification. PLoS ONE, 7(11), e49790. 10.1371/journal.pone.0049790

[8]. Chandio, B. Q., Risacher, S. L., Pestilli, F., et al. (2020). Bundle analytics, a computational framework for investigating the shapes and profiles of brain pathways across populations. Scientific Reports, 10, 17149. 10.1038/s41598-020-74054-4

[9]. I. Ba Gari, S. Javid, A. Ramesh, N. Jahanshad, et al., “Along-tract parameterization of white matter microstructure using medial tractography analysis (MeTA),” Proc. SPIE Medical Imaging/SIPAIM, 2023. 10.1109/SIPAIM56729.2023.10373540

[10]. Yendiki, A., et al. (2011). Automated probabilistic reconstruction of white-matter pathways in health and disease using an atlas of the underlying anatomy. Frontiers in Neuroinformatics, 5, 23. 10.3389/fninf.2011.00023

[11]. Wasserthal, J., Neher, P., & Maier-Hein, K. H. (2018). TractSeg: Fast and accurate white matter tract segmentation. NeuroImage, 183, 239–253. 10.1016/j.neuroimage.2018.07.070

[12]. Chamberland, M., St-Jean, S., Jones, D. K., Descoteaux, M., & Leemans, A. (2025). Methods and statistics for diffusion MRI tractometry. Handbook of Diffusion MR Tractography, 439–450. Academic Press. 10.1016/B978-0-12-818894-1.00023-9

[13]. Everitt, B. J., & Robbins, T. W. (2016). Drug addiction: Updating actions to habits to compulsions ten years on. In Neuroscience of Addiction (Elsevier). 10.1016/B978-0-12-818894-1.00023-9

[14]. Cai, L. Y., Yang, Q., Hansen, C. B., et al. (2021). PreQual: An automated pipeline for integrated preprocessing and quality assurance of diffusion MRI images. Magnetic Resonance in Medicine, 86(1), 456–470. 10.1002/mrm.28678

[15]. Feng, Y., Shuai, Y., Villalón-Reina, J. E., Chandio, B. Q., Thomopoulos, S. I., Nir, T. M., Jahanshad, N., & Thompson, P. M. (2026). Streamline density normalization: A robust approach to mitigate bundle variability in multi-site diffusion MRI. In Computational Diffusion MRI (CDMRI 2025), Lecture Notes in Computer Science, 16205, 44–56. Springer, Cham. 10.1007/978-3-032-12837-9_5

[16]. Tournier, J.-D., Smith, R., Raffelt, D., et al. (2019). MRtrix3: A fast, flexible and open software framework for medical image processing and visualisation. NeuroImage, 202, 116137. 10.1016/j.neuroimage.2019.116137

[17]. Avants, B. B., Tustison, N. J., Song, G., Cook, P. A., Klein, A., & Gee, J. C. (2011). A reproducible evaluation of ANTs similarity metric performance in brain image registration. NeuroImage, 54(3), 2033–2044. 10.1016/j.neuroimage.2010.09.025

[18]. St-Onge, E., Garyfallidis, E., & Collins, D. L. (2022). Fast streamline search: An exact technique for diffusion MRI tractography. Neuroinformatics, 20(4), 1093–1104. 10.1007/s12021-022-09590-7

[19]. Chandio, B. Q., et al. (2024). Bundle analytics–based data harmonization for multi-site diffusion MRI tractometry. Proceedings of the 46th Annual International Conference of the IEEE Engineering in Medicine and Biology Society (EMBC), Orlando, FL, USA, pp. 1–7. 10.1109/EMBC53108.2024.10782419

[20]. Ba Gari, I., Bhatt, R. R., Yeh, F.-C., & Jahanshad, N. (2025). Heritability and genetic correlations along the corticospinal tract. In Computational Diffusion MRI (CDMRI 2024), Lecture Notes in Computer Science, 15171. Springer, Cham. 10.1007/978-3-031-86920-4_18

[21]. Benjamini, Y., & Hochberg, Y. (1995). Controlling the false discovery rate: a practical and powerful approach to multiple testing. Journal of the Royal Statistical Society: Series B, 57(1), 289–300.10.1111/j.2517-6161.1995.tb02031

[22]. Feng, Y., Villalón-Reina, J. E., Ba Gari, I., Alibrando, J. D., Nir, T. M., Jahanshad, N., Chandio, B. Q., & Thompson, P. M. (2025). Evaluating sample-size efficiency and sensitivity of tractometry in Alzheimer’s disease. medRxiv (preprint). 10.1101/2025.11.11.687878

[23]. Song, S.-K., Sun, S.-W., Ramsbottom, M. J., Chang, C., Russell, J., & Cross, A. H. (2002). Dysmyelination revealed through MRI as increased radial (but unchanged axial) diffusion of water. NeuroImage, 17(3), 1429–1436. 10.1006/nimg.2002.1267

[24]. Song, S.-K., Yoshino, J., Le, T. Q., et al. (2005). Demyelination increases radial diffusivity in corpus callosum of mouse brain. NeuroImage, 26(1), 132–140. 10.1016/j.neuroimage.2005.01.028

[25]. Beaulieu, C. (2002). The basis of anisotropic water diffusion in the nervous system –a technical review. NMR in Biomedicine, 15(7–8), 435–455. 10.1002/nbm.782

